# Multilevel sex-specific neurobiological signatures of early life adversity

**DOI:** 10.64898/2026.01.26.701291

**Authors:** Sowmya Narayan, Christina Beer, Carlo Castoldi, Tibor Stark, Beatrice Dal Bianco, Simone Röh, Susann Sauer, Joeri Bordes, Stoyo Karamihalev, Shiladitya Mitra, Veronika Kovarova, Patricia Maidana Miguel, Bonnie Alberry, Darina Czamara, Patricia Pelufo Silveira, Michael Czisch, Bianca Silva, Elisabeth B. Binder, Mathias V. Schmidt

## Abstract

Stress exposure early in life is an established risk factor for adult psychiatric illness, yet these disorders – including anxiety disorders and depression – show significant sex-dependence in prevalence, symptomatology, and treatment response. The biology underlying these differences remains largely unexplored and may contribute to the clinical heterogeneity in anxiety and depression. Here, we characterize the lasting impact of developmental stress on adulthood neurobiology and behavior in mice by combining analyses of multiple levels of brain function, including whole-brain c-Fos mapping, manganese-enhanced MRI and transcriptomics with advanced behavioral phenotyping. Across levels of investigation, we find distinct and often opposite effects of developmental stress depending on sex. These results together showcase the strong influence of sex on how early life adversity affects the onset of stress-related disorders. This work emphasizes the necessity of considering sex when investigating developmental and neurobiological underpinnings of stress-related disorders and displays a vast range of lasting effects of developmental stress on the brain, which provides a valuable resource for future studies aiming to improve psychiatric treatments.

## Main

Stress exposure during neurodevelopmental time periods is an established risk factor for psychiatric disorders, such as depression and anxiety disorders^1–3^. Exposure to prenatal and early life stress are unfortunately commonplace, and psychopathology is seen to be more challenging and resistant to treatment when it is associated with developmental exposure to adversity^1,4^. While it is known that stress during these time periods incites biological changes that directly affect the developing brain and influence later psychopathology, the specific neurobiological mechanisms that link these developmental stressors to adult psychopathology remain to be fully understood.

A critical, yet often neglected, factor is sex. There are well-documented sex differences in the incidence of stress-related disorders, with women and sexual- and gender-minority groups often experiencing higher rates of adversity and psychiatric illness^5–8^. The brain exhibits natural sexual differentiation during developmental processes in regions which have important functional roles in processing stress, such as the amygdala, hippocampus, hypothalamus and BNST ^9,10^, and there is evidence of differing interactions of male and female gonadal hormones with the sustained hormonal stress response, the hypothalamic-pituitary-adrenal (HPA) axis^11,12^. To this end, many studies show sexually divergent effects of developmental stressors in the brain, which converge onto behavioral phenotypes^13–27^. However, a holistic characterization of the lasting, sex-specific effects of developmental stress on the brain and behavior is still needed to fully understand risk mechanisms for mental illness.

To address this, we developed a translational mouse model which combines prenatal and early life stress exposures (PNELS) to conduct a comprehensive sex-specific analysis of developmental stress. By combining automated deep learning-based behavioral phenotyping with multi-level neurobiological profiling – including sustained neural activity, functional circuit activation, and transcriptomics (see Fig. 1A for summary of study design) – we uncover distinct, sex-specific pathways by which early adversity impacts adult behavior and brain function. We find that sex strongly shapes neurobiological and behavioral responses to developmental stress, and we identify specific molecular and circuit-level signatures associated with a greater risk of depression-like behavior in adulthood.

**Figure 1.**
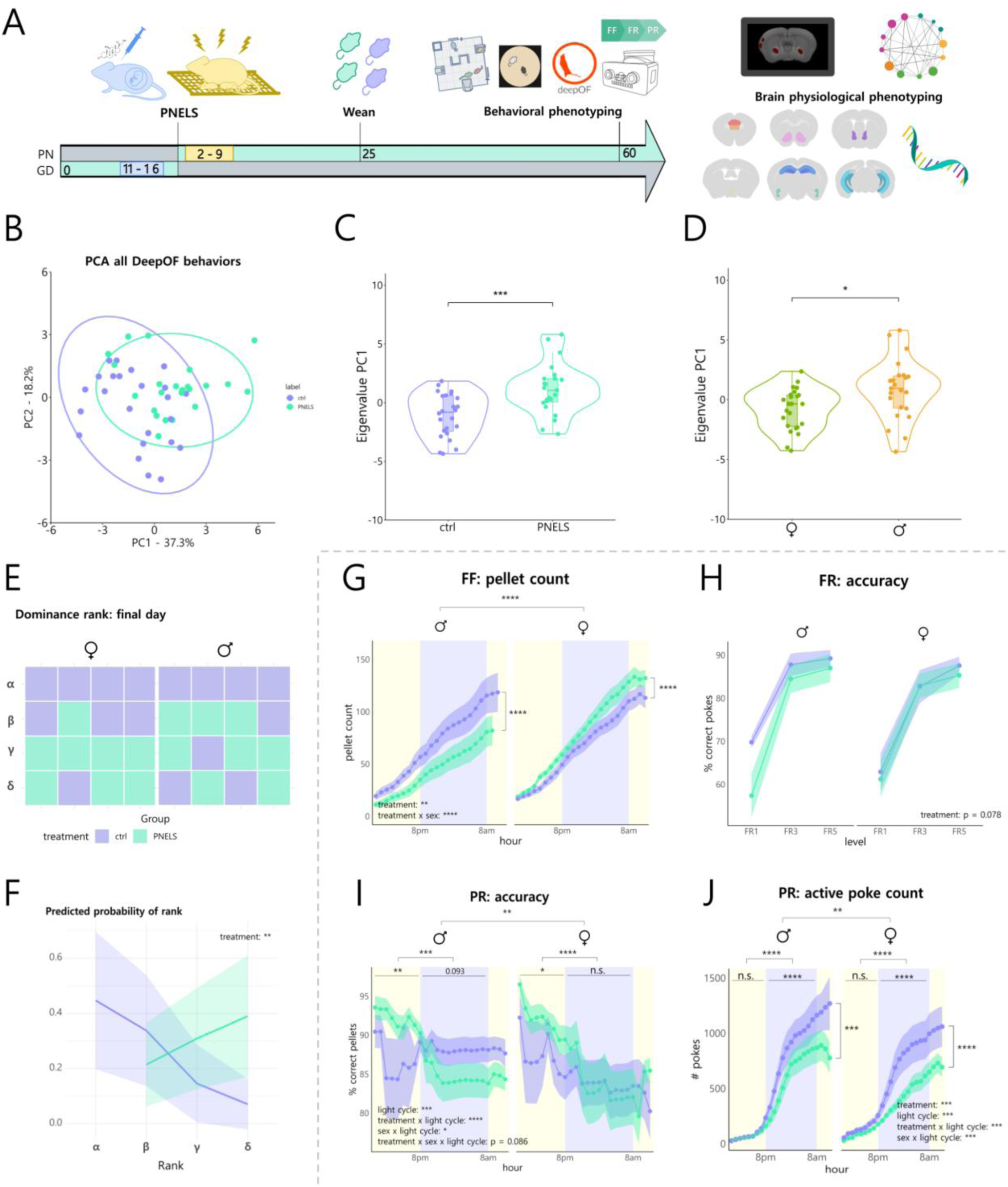
Developmental stress leads to alterations in adulthood behaviors related to psychiatric disorders. Timeline of PNELS experiments; all animals were first exposed to prenatal stress (CORT injection during GD 11-16) and early life stress (limited bedding and nesting during PN 2-9) (**A**). In the OFSI test, PCA of all DeepOF behaviors (**B**) shows differences between treatment groups (**C**, Wilcoxon test: W = 124, p = 0.000925) and sexes (D, Wilcoxon test: W = 157, p = 0.0107). In the semi-naturalistic social box paradigm, PNELS mice were always in subordinate positions in the social dominance hierarchy (**E**), making PNELS treatment predictive of rank (**F**, estimate = −2.1281, SE = 0.7519, z = −2.83, p = 0.00465). In the FED3 paradigm, free feeding pellet differed between sexes in the effect of PNELS (**G**, ANOVA: treatment: F(1, 992) = 9.151, p = 0.0026; sex: F(1, 992) = 72.139, p < 0.0001; time: F(23, 992) = 50.474, p < 0.0001; treatment x sex: F(1, 992) = 95.751, p < 0.0001, Post-hoc Tukey: M ctrl vs PNELS: p < 0.0001; F ctrl vs PNELS: p < 0.0001), while percentage of active pokes in the fixed ratio stages was similar between sexes with only a trend towards lower accuracy in the PNELS group (**H**, ANOVA: treatment: F(1, 114) =3.156, p = 0.0783; Post-hoc Tukey: all ns). Task accuracy is higher in all PNELS animals during the daytime but switches at nighttime to reduced accuracy in PNELS males and no differences in females (**I**, ANOVA: treatment: F(1, 956) = 3.272, p = 0.0708; sex: F(1, 956) = 5.145, p = 0.0235; light cycle: F(1, 956) = 54.396, p < 0.0001; treatment x light cycle: F(1, 956) = 25.902, p < 0.0001; sex x light cycle: F(1, 956) = 5.851, p = 0.0158; treatment x sex x light cycle: F(1, 956) = 2.946, p = 0.0864. Post-hoc Tukey: day ctrl vs PNELS: p < 0.0001, night ctrl vs PNELS: p = 0.3128; day M vs F: p = 0.9573, night M vs F: p = 0.0062). Amount of active pokes is lower in PNELS animals for both sexes (**J**, ANOVA: treatment: F(1, 958) = 53.668, p < 0.0001; sex: F(1, 958) = 9.185, p = 0.0025; light cycle: F(1, 958) = 538.219, p < 0.0001; treatment x light cycle: F(1, 958) = 23.096, p < 0.0001; sex x light cycle: F(1, 958) = 13.587, p = 0.0002. Post-hoc Tukey: F ctrl vs PNELS: p < 0.0001, M ctrl vs PNELS: p = 0.0003; day ctrl vs PNELS: p = 0.8807, night ctrl vs PNELS: p < 0.0001; day M vs F: p = 0.7325, night M vs F: p < 0.0001). Stars represent p-values as follows: * < 0.05, ** < 0.01, *** < 0.001, **** < 0.0001.

With this study, we provide an expansive foundational resource for a more precise, sex-informed approach to understanding and treating psychiatric illness, and emphasize the critical need to consider sex as a biological variable in both preclinical research and the development of targeted therapies for stress-related disorders.

## Results

### Developmental stress leads to alterations in adulthood behaviors related to psychiatric disorders

We first assessed whether developmental stress leads to enduring, sex-specific alterations in behaviors relevant to psychiatric disorders. To achieve this, we employed a deep behavioral phenotyping pipeline that utilized marker-less pose estimation (DeepLabCut^28^) and a supervised behavioral annotation framework (DeepOF^29,30^)(Fig. 1A, Supplementary Fig. 1A-C). An open field and social interaction (OFSI) test^29^ with freely interacting mice was used to investigate social and anxiety-like behaviors. The first principal component (PC1) of a principal component analysis (PCA), used to represent overall variation in these stress-related behaviors (Fig. 1B), showed a difference due to developmental stress (PNELS) (Fig. 1C; W = 124, p = 0.0009) and between sexes (Fig 1D; W = 157, p = 0.0107).). To further characterize the effects of PNELS on social behavior, we assessed social dominance using the social box paradigm^31^, where both male and female PNELS animals adopted exclusively subordinate positions of the social dominance hierarchy (Fig. 1E). Developmental stress (PNELS) was found to be a significant predictor of rank in the hierarchy (Fig. 1F, p = 0.0047).

In a sequential set of home-cage paradigms using sucrose pellet rewards to assess depression-related behaviors^32^ (Supplementary Fig. 1C n = 12 for each group), PNELS males displayed reduced passive reward consumption (p < 0.0001), consistent with classical reports of anhedonia-like behavior. In contrast, PNELS females consumed more than controls (p < 0.0001), suggesting divergent effects of stress on reward sensitivity between sexes (Fig. 1G). In the fixed ratio task stages^32^ FR1, FR3, and FR5 (Supplementary Fig. 1C), task accuracy did not differ due to PNELS or sex, but PNELS animals showed a trend towards lower task accuracy which was driven by the male FR1 stage (Fig. 1H; p = 0.0780), suggesting possible PNELS-linked learning impairments in males alone. In the progressive ratio (PR) task, PNELS animals showed higher accuracy during daytime which decreased at night—more so in males, with no difference in females (Fig. 1I). Active pokes followed a similar pattern, with PNELS animals exhibiting lower activity which was most significant during nighttime (Fig. 1J, p < 0.0001). In males particularly, reduced night time accuracy despite sustained activity suggests increased impulsivity due to PNELS.

Together, these findings support that developmental stress has long-term, sex-specific effects on depression- and anxiety-like behaviors. Our assessments uncovered robust, sex-dependent behavioral patterns, which are often missed in classical tests (Supplementary Fig. 2). In particular, specific behavioral domains including social, anxiety-like, and reward-motivated behaviors are affected by developmental stress sex-specifically, which are often ignored in standard testing.

### Baseline brain activity in regions relevant for stress processing is altered in adulthood in a sex-dependent manner

To understand brain-wide sustained activity differences in adulthood induced by developmental stress, we employed manganese-enhanced MRI (MEMRI) following sustained Mn²⁺ uptake via osmotic pump infusion (Supplementary Fig. 1C; ctrl M: n=9; PNELS M: n=8; ctrl F: n=12; PNELS F: n=11). Here, MEMRI contrast differences reflect activity dependent accumulation of Mn2+ averaged over a whole week (p_FDR,cluster_ < 0.05).

Voxel-wise analysis revealed a predominant main effect of PNELS in increasing Mn^2+^-enhanced contrast mainly in Superior Colliculus, Entorhinal areas, Lateral Septum, Medial Preoptic Area, cortices (primary somatosensory, Posterolateral visual area, Agranular Insula posterior part), Endopiriform nucleus ventral part, and central amygdala (Fig. 2A, sex: pink, PNELS: turquoise, interaction: yellow; Supplementary Fig. 3), as well as wide range of sex-specific stress effects in a number of different brain regions including cortical, subcortical and brainstem areas, with mainly intensity increases for stressed males and decreases for stressed females (Fig. 2B, 2C). Common stress-related regions, including the amygdala, bed nucleus of the stria terminalis (BNST), nucleus accumbens (NAc), and hypothalamic, thalamic, and cortical structures, were found in clusters with significant interaction effects and in pairwise comparisons between groups, indicating broad sex-specific effects of PNELS on sustained brain activity that mirror the distinct behavioral patterns.

**Figure 2.**
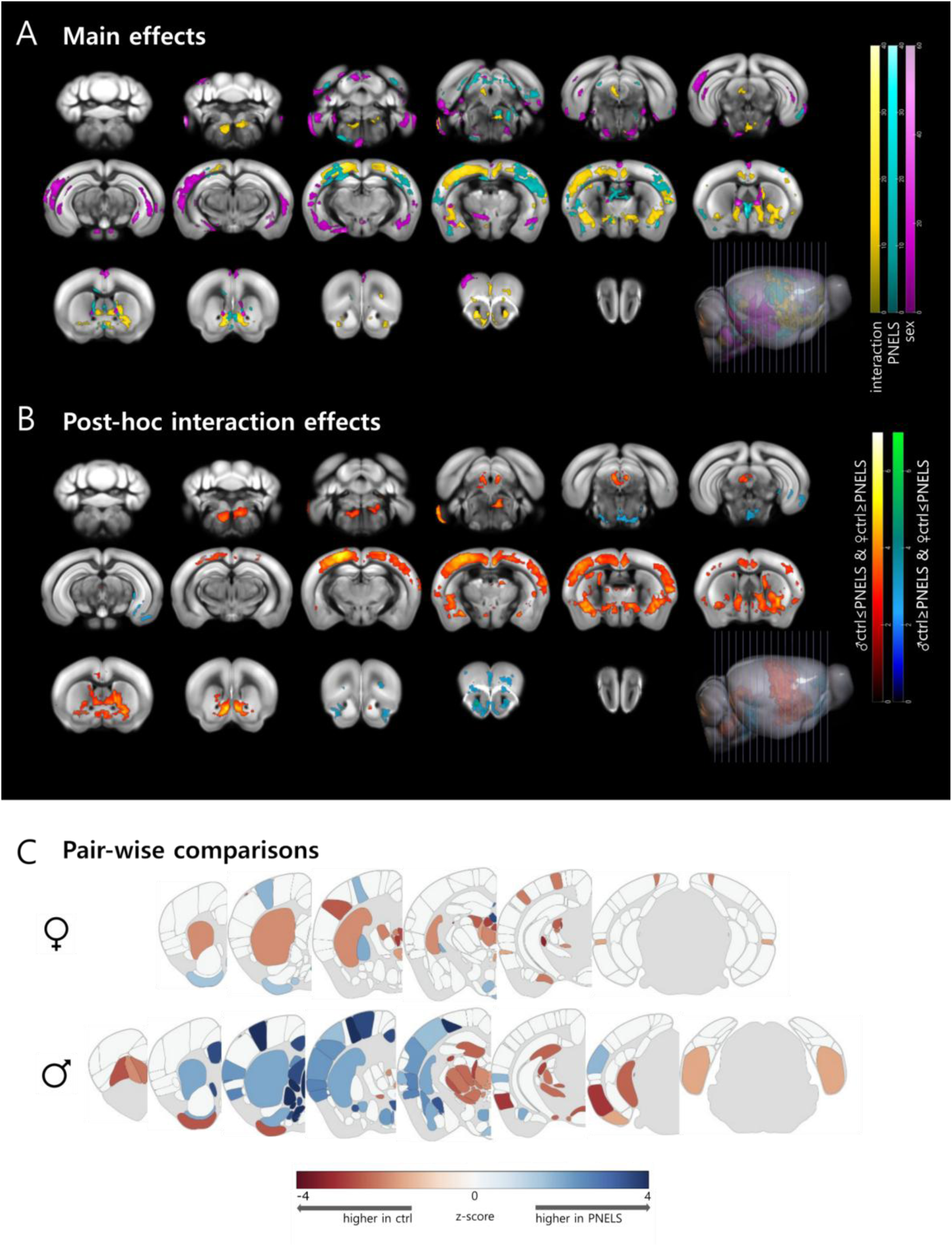
Baseline brain activity in regions relevant for stress processing are altered in adulthood in a sex-dependent manner. Main effects (**A**) of PNELS, sex, and their interaction in brain activity evaluated by voxel-wise analysis of MEMRI found significant (F-test, p_FDR,cluster_ < 0.05) changes in common stress related regions including amygdala, BNST, nucleus accumbens (NAc), hypothalamus, thalamic structures, and cortex (**B**). Pairwise comparisons of voxel intensities between ctrl and PNELS groups of each sex indicate opposing direction of effect of PNELS in stress related regions (t-test, p_FDR,cluster_ < 0.05) (**C**). Statistical maps are presented with clusters surviving FDR cluster correction (pFDR,cluster < 0.05), using a collection threshold of uncorrected p < 0.005.

### Developmental stress has sex-specific effects on brain functional networks underlying behaviors relevant for psychiatric disorders

To further probe the neural correlates of behavioral alterations induced by developmental stress, we next performed whole-brain c-Fos mapping following the OFSI paradigm (Supplementary Fig 1B, n = 8 for each group and sex). We again observed sex-specific effects of developmental stress on global c-Fos expression. A region-by region analysis revealed that in males, PNELS increased c-Fos expression relative to the control condition in regions where control females showed higher baseline activity, such as in thalamic, hypothalamic, and some isocortical areas (Fig. 3A, B) suggesting sex-specific activity reorganization following stress during early development. Functional connectome analyses further supported this dissociation. Although PNELS males showed a less prominent global activity shift, they showed a marked reduction in interregional connectivity relative to controls, whereas PNELS females exhibited the opposite pattern, with enhanced connectivity strength and density (Fig. 3C, R ≥ 0.87; p ≤ 0.0025).

**Figure 3.**
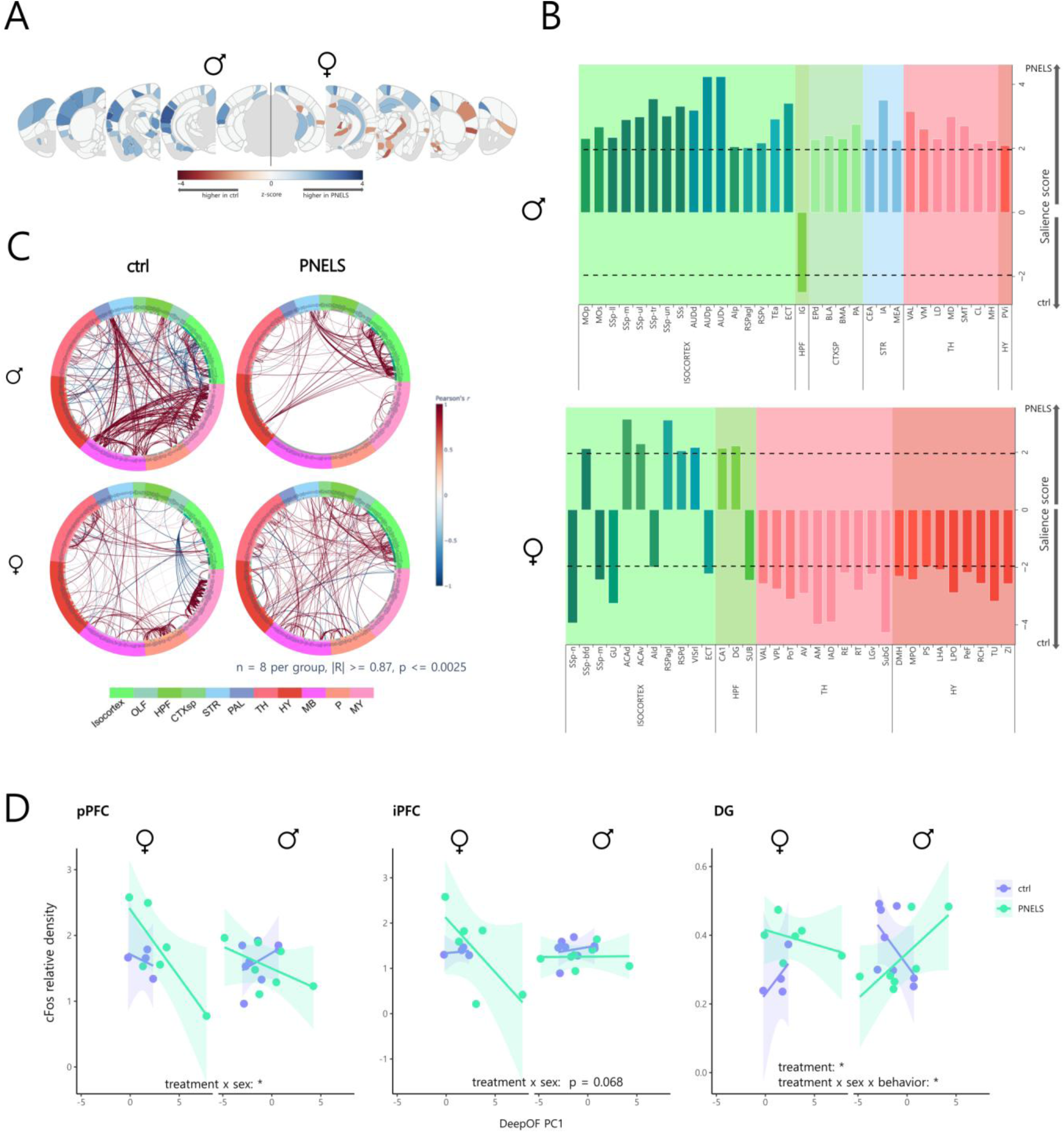
Developmental stress has sex-specific effects on brain functional networks underlying behaviors relevant for psychiatric disorders. Direction of contribution of cFos activity to stress in brain subregions, in males (global p-value=0.227) and females (global p-value=0.075)(**A**, **B**; p < 0.05) and functional connectome of each group (**C** R≥0.87, p<0.0025) showed that males and females showed many opposing patterns of brain network connectivity in adulthood after PNELS. Linear regression to show whether cFos relative density values from select brain regions, sex, and DeepOF behavioral composite score can predict cFos relative density (**D,** lm(region ∼ sex * treatment * DeepOF PC1); pPFC: treatmentPNELS, p = 0.073, sexM:treatmentPNELS, p = 0.0416; iPFC: treatmentPNELS, p = 0.092, sexM:treatmentPNELS: p = 0.068; vDG: treatmentPNELS: p = 0.03723, sexM:treatmentPNELS:PC1: p = 0.031)

To identify associations between the behavioral and brain network alterations following PNELS, we focused on regions relevant to stress processing and used linear regression to associate c-Fos relative density values with a behavioral composite score (PC1) calculated from the open field and social interaction (OFSI) test. The prelimbic and infralimbic cortex and the dentate gyrus stood out as regions where predictive relationships between c-Fos activity and PC1 behavioral scores were altered by both sex and developmental stress (Fig. 3D, p < 0.05).

Together, these results reveal that developmental stress reshapes circuit activation and functional connectivity in a sex-specific manner, with implications for divergent behavioral outcomes.

### Diverse changes in the brain transcriptomic responses indicate sex-specific effects of developmental stress

To investigate the molecular basis of these differential brain activity and behavioral patterns, we next performed bulk RNA sequencing in 10 selected regions of interest (Supplementary Fig. 1A), selected for their known roles in stress-related psychopathology. Differential gene expression analysis from bulk RNA-sequencing revealed region- and sex-specific transcriptional responses to PNELS (FDR-adjusted p (padj) < 0.1). The majority of differentially expressed genes (DEGs, 93.7% in males and 98.2% in females) were unique to a single region, and the amount of DEGs per region varied markedly by sex (Fig. 4A; Supplementary Table 1) with the highest number of DEGs in males identified in the NAc (N=67) and the ventral CA1 (N=60) in females. These findings underscore the distinct biological roles and transcriptional architectures of each brain region in the context of developmental stress.

**Figure 4.**
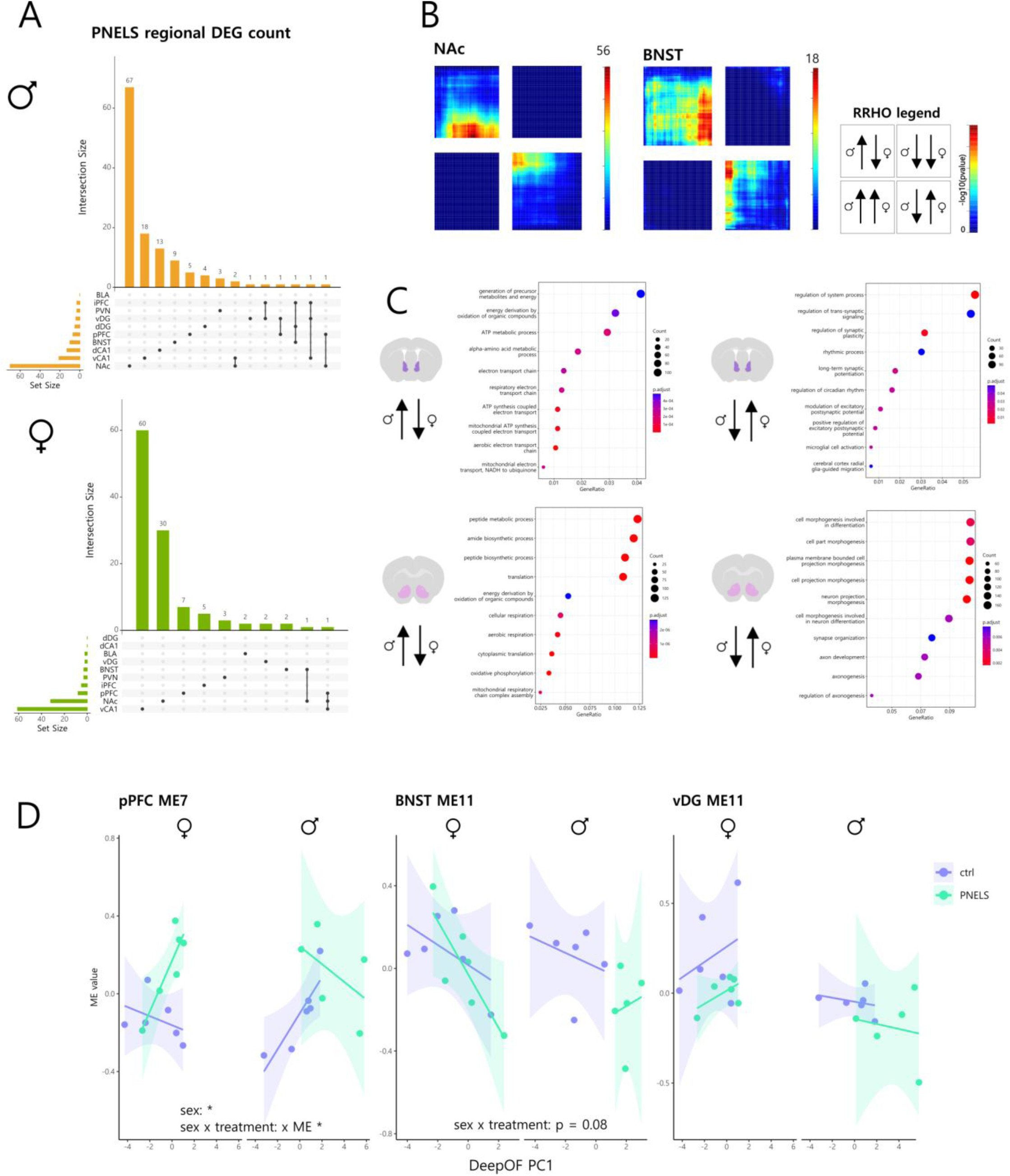
Diverse changes in the brain transcriptomic responses indicate sex-specific effects of developmental stress. The majority genes differentially expressed (p <0.1) due to PNELS were not shared across brain regions (**A**). Threshold-free overlap of commonly expressed male and female gene lists found discordance in NAc and BNST (**B**). Gene ontology enrichment for the lists discordant between sexes indicate opposite direction of regulation of neuronal signaling and cell metabolic process (**C**). WGCNA revealed modules strongly associated to PNELS, sex, and DeepOF PC1 (R>0.5; p<0.05) of which pPFC ME7 and BNST ME 11 eigenvalues had predictive relationships with behavior which were dependent on sex and treatment (**D,** lm(DeepOF PC1 ∼ sex * treatment * ME eigenvalues); pPFC ME7: sexM, p = 0.0379, sexM:treatmentPNELS:ME7, p = 0.0102; BNST ME11: sexM:treatmentPNELS, p = 0.08; vDG ME11: all ns).

As very few DEGs were shared between sexes at the FDR cut-off (FET p > 0.5), we applied a threshold-free rank–rank hypergeometric overlap (RRHO) analysis to examine transcriptomic convergence or divergence between sexes. While most regions displayed largely concordant expression changes across sexes after PNELS (Supplementary Fig. 4A), the nucleus accumbens (NAc) and bed nucleus of the stria terminalis (BNST) showed divergence in direction of gene regulation between sexes with mostly opposite effects observed (Fig. 4B; ρ < 0, CI < 0.5). These regions, implicated in reward processing and anxiety, may serve as sex-specific hubs mediating observed behavioral responses to developmental stress.

Functional enrichment of genes regulated in opposite directions between sexes in the NAc and BNST (Supplementary Table 2) further supported this dissociation. In males, upregulated pathways included energy metabolism and cellular homeostasis, while in females, opposing changes were enriched for synaptic signaling, plasticity, and neurodevelopmental processes (Fig. 4C).

To identify coordinated transcriptional networks associated with PNELS, we performed weighted gene co-expression network analysis (WGCNA) in regions of interest where previous data implicated sexual dimorphism in developmental stress outcomes (Supplementary Fig. 5). Three modules in the pPFC, BNST, and vDG showed significant correlations and predictive relationships with sex and stress exposure (Fig. 4D, p < 0.05) revealing candidate gene networks that may underlie sex-specific vulnerability to developmental stress.

### Sex differences in mechanisms of developmental stress contributing to major depressive disorder in humans

To assess the translational relevance of transcriptomic alterations observed in the PNELS model, we compared our mouse data to a publicly available human dataset profiling gene expression in the post-mortem pPFC, iPFC, and NAc of individuals with major depressive disorder (MDD)^33^. Interestingly, though concordance was low between species (ρ ≈ 0, CI ≈ 0.5), we observed opposing patterns of direction of gene regulation between sexes across each region (Fig. 5A), suggesting that developmental stress may engage distinct transcriptional programs in males and females that converge on divergent risk trajectories for depression.

**Figure 5.**
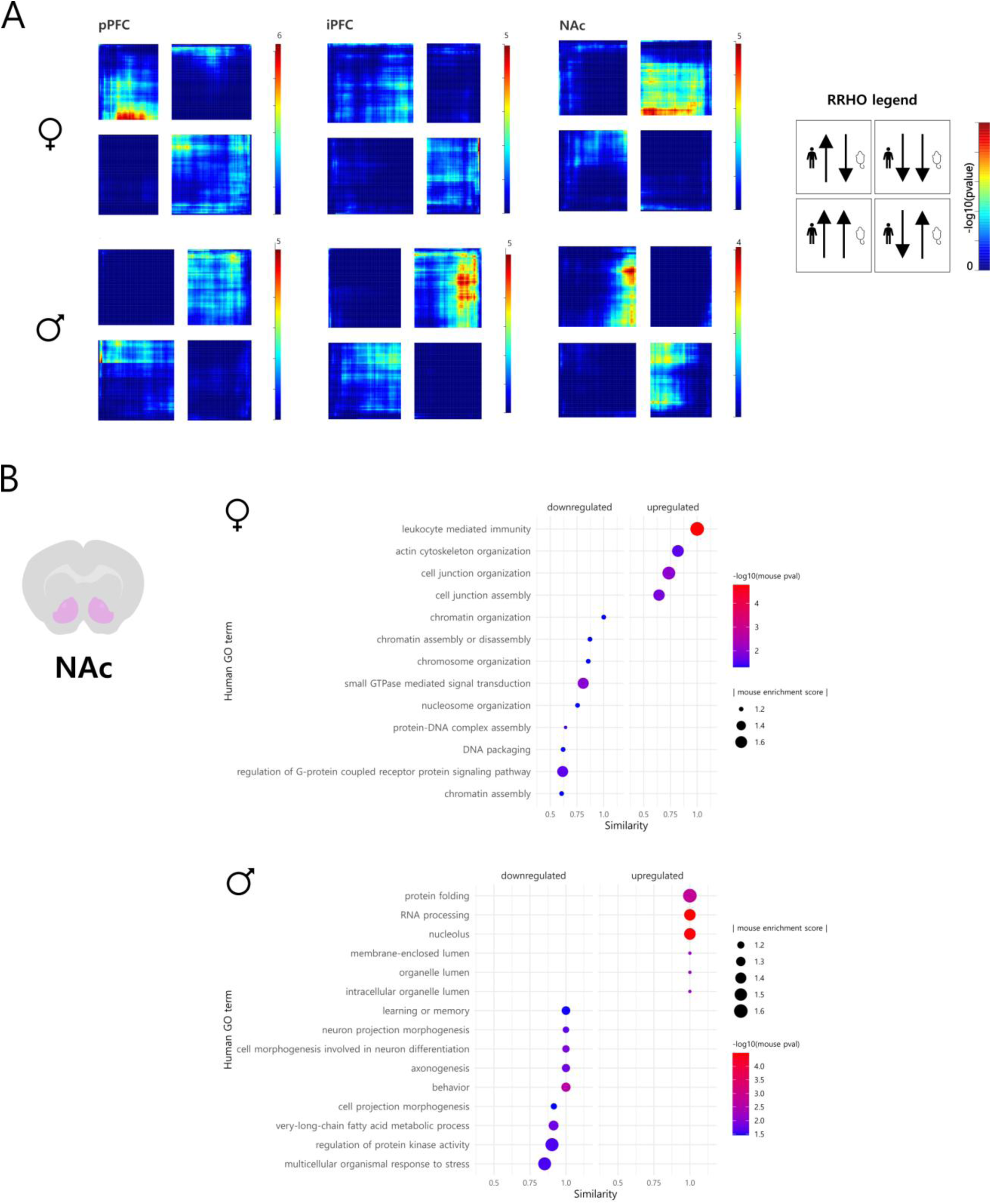
Sex differences in mechanisms of developmental stress contributing to depression onset. RRHO for male and female MDD profiles against PNELS profiles in pPFC, iPFC, and NAc showed an opposite pattern of direction of regulation between sexes in overlapping genes per region (**A**). Semantic similarity analysis indicates sex specific terms overlapping across PNELS in mice and MDD in humans (**B**, p<0.05, similarity>0.5), with processes related to cellular and DNA organization more conserved in females and processes more related to neuronal function in males.

Furthermore, using available human data for the NAc, we assessed semantic similarity of gene ontology enrichment for developmental stress and major depressive disorder, finding distinct biological processes between species for male and female lists (Supplementary Table 4). Males had more overlap between species and upregulated terms related to nuclear processes downregulated to neuronal and behavioral terms, while females had upregulated terms related to cellular assembly and downregulated terms related to DNA organization (Fig. 5B).

These findings underscore the importance of considering sex-specific molecular pathways in the etiology of stress-related disorders and suggest that developmental stress can prime the brain for depression through distinct, region- and sex-dependent mechanisms.

## Discussion

In this study, we characterized sex-specific effects of developmental stress exposure on adulthood behavior and neurobiology, in a novel, translationally relevant mouse model. By combining deep behavioral phenotyping with multimodal investigation, including baseline brain-wide activity, circuit-level function, and bulk transcriptomic profiling, we offer a comprehensive view of how developmental stress shapes the brain and behavior across multiple levels of organization in both sexes.

A central finding of our work is the profound sexual dimorphism in how early life adversity impacts the brain and behavior. This observation builds on and significantly extends previous literature documenting substantial sex differences in response to stress and adversity^14,34–36^. Our data reveal convergent and distinct effects of developmental stress across behavioral domains, which could largely explain the sex-dependent prevalence and symptomatology of stress-related psychopathology reported in humans^22,37^. This work challenges classical, male-centric interpretations of stress-related behavior and indicates the importance of including sex-specific investigations in stress and psychiatric research. It offers new insight into how stress responses may reflect distinct adaptive or maladaptive strategies in males and females.

At the neurobiological level, we found that PNELS induces widespread, sex-dependent alterations in brain activity patterns. In the MEMRI data, PNELS recruits a coherent network for stress response, with heightened basal activity in regions involved in hypervigilance, anticipatory anxiety, and autonomic tone. Interaction effects between sex and PNELS are showing mainly increases for males after PNELS, of which some activity difference may be linked to the observed behavioral differences, such as higher impulsivity in the sucrose reward task to elevated hypothalamic (arousal) and NAc activity in stressed male mice^38^.

Functional c-Fos mapping of task-evoked activity revealed different and often opposing network-level strategies for stress adaptation in males versus females, potentially speaking to the interplay of gonadal hormones on modulating neuroplasticity, connectivity, and neurotransmitter signaling^39^. In males, predominantly cortical and limbic brain areas showed hyperactivation following early adversity, while in females thalamic and hypothalamic nuclei displayed hypoactivation. Collectively, these findings suggest that even similar behavioral consequences of early life adversity, such as lower social dominance^40^ and suppressed reward behavior, are governed by sex-distinct changes in brain circuit activity.

Our transcriptomic analysis further supports this divergence, revealing near-opposite directions of gene regulation due to developmental stress in key brain regions—particularly the nucleus accumbens (NAc) and bed nucleus of the stria terminalis (BNST). These regions are known to mediate reward and anxiety-related behaviors and are strongly interconnected^41,42^. Notably, enrichment analyses suggest that PNELS affects synaptic signaling, neurodevelopment, and metabolic pathways in a sex-specific manner. Our data suggest that sex differences in molecular response to early stress, particularly in regions related to reward-processing and anxiety, may play a central role in shaping long-term vulnerability or resilience to psychiatric disorders.

These neurobiological insights were further strengthened by a cross-species comparison using post-mortem human brain data from individuals with major depressive disorder (MDD). Our findings in mice with PNELS are similar to sex-dimorphic findings of transcriptional changes in postmortem brain of individuals suffering from MDD^33^, highlighting the importance of sex-specific effects. When comparing these signatures across species, we observed only slight overlap, but in a sex-specific manner, concordant or discordant between sexes depending on the region, suggesting that MDD transcriptomic profiles reflect changes including but also going beyond stress-related effects. Enriched biological processes between PNELS and MDD were also distinct between sexes, with implications for the development of targeted, sex-sensitive interventions.

In conclusion, this study’s translational approach, multimodal neurobiological analysis, and rigorous sex-specific investigation reveal novel insights into the lasting consequences of early life adversity. By laying a foundation for more inclusive and precise models of psychiatric vulnerability, this work encourages more effective and equitable strategies for prevention and treatment.

## Methods

### Mouse model

#### Housing conditions

All protocols were approved by the committee for the Care and Use of Laboratory Animals of the government of Upper Bavaria, Germany and all procedures were in accordance with the European Communities Council directive 2010/63/UE.

Adult CD1 mice aged 8-10 weeks were obtained from Charles River and housed in standard conditions (12h/12h light/dark cycle with dark phase starting at 8pm, temperature 23 ± 2°C, 55% humidity) at the Max Planck Institute for Psychiatry, with laboratory chow diet and water available *ad libitum*. Animals were acclimated for one week in the facility before beginning procedures.

#### Breeding

Male and female mice were pair-housed and females checked daily for vaginal plugs to estimate date of conception. Males and females were separated after 5 days of breeding.

#### Developmental stress treatment

A combination of prenatal and postnatal early life stressors was used, to increase translational value.

The protocol for prenatal stress exposure (PNS) was derived from Astiz and colleagues^43^. Solid 98% corticosterone (CORT) was dissolved in polyethylene glycol-400 (*Sigma-Aldrich*) at a concentration of 20mg/ml and stored at room temperature. Pregnant dams were injected subcutaneously with CORT daily at a concentration of 50mg/ml from gestational day (GD) 11 through GD16 within the first hour of the lights turning on in the animal facility, to offset natural corticosterone rhythms.

The established limited bedding and nesting (LBN) paradigm^44^ was introduced as an early life stressor (ELS). On postnatal day 2 (D2), litters were culled to a maximum of 12 pups, balancing as much as possible for sex. Dams and pups were transferred to a cage a metal grid inlayed on the floor, which contained half of a cotton nestlet and 50ml of bedding material. Control litters were transferred to a cage with the standard amount of bedding material and 2 cotton nestlets. At D9, pups were weighed and transferred back to standard housing conditions.

Each experimental cohort included two experimental groups: non-stressed controls (ctrl) and a combination of prenatal and early life stress (PNELS) with an equal number of animals of each sex in each group.

#### Weaning

Between postnatal days 24-26, animals which were to be used for social behavioral phenotyping were weaned into non-sibling social groups containing two PNELS animals and 2 ctrl animals. Animals were weighed on the weaning day and all behavioral phenotyping began in young adulthood, from 8 weeks of age.

### Behavioral phenotyping

#### Open field + social interaction (OFSI) test

Open field and social interaction tasks were performed in an acute, novel context as previously described^29^. Experimental animals were placed in a round open field (diameter 38cm) with bedding material on the apparatus floor. Exploration activity of the single animal was recorded for 10 minutes, then an unfamiliar C57BL6 of similar age was introduced into the apparatus as a novel social partner and pair activity was recorded for another 10 minutes.

#### Social box

The social box was used to test group social dynamics and dominance hierarchy in a semi-naturalistic setting. The protocol and setup were implemented as previously published^31^. Mice were marked with distinguishable colors and placed in groups of four in a 60 x 60cm square arena, where their activity was recorded for 4 days and 4 nights.

#### FED3 home-cage operant reward task

Feeding Experiment Devices version 3.0^32^ (FED3) were used to assess behaviors related to motivation, anhedonia, and reward learning. Each FED3 has a programmed active and inactive nose port and a mechanism to contain and dispel 20mg sucrose pellets (*5TUT*). Mice were first habituated with single housing and 15 sucrose pellets scattered throughout the home cage. On the first experimental day, a FED3 was added to the home cage in free feeding (FF) mode, in which sucrose pellets were readily retrievable. Animals were checked once daily and progressed to the fixed-ratio (FR) stages once they retrieved a minimum of 100 pellets in the FF stage. For the stages FR1, FR3, and FR5, the animal needed to perform one, three, or five nose-pokes, respectively, in the active port in order to receive a sucrose pellet. Animals were progressed fromFR1 to FR3 after receiving at least 75 pellets in FR1, and to FR5 after 60 pellets in FR3. After 60 pellets in FR5, animals were food-restricted for 24 hours and proceeded to the final progressive ration (PR) stage. During the PR, the number of nose pokes required to receive a reward increased after each reward was retrieved. Analysis of FED3 data was implemented with the aid of python package FED3_Viz-0.5.1 and R package FED3.

### Manganese-enhanced MRI (MEMRI)

#### Subject preparation

Animals were sedated with 4% isoflurane, secured in a prone stance on a platform, and maintained at 1.5-2.5% isoflurane anesthesia (100% oxygen, airflow 1.5 liter/min) throughout the scanning procedure. Bepanthen® cream was applied to the eyes to prevent drying. Body temperature was monitored via a rectal temperature and maintained at 36-38°C, and respiration was monitored with a pressure sensor set under the animal’s thorax. Isoflurane flow was adjusted accordingly as needed to maintain a respiration rate of 80 - 120 breaths per minute. Animals were sacrificed after the scanning procedure.

#### Manganese infusion

Manganese(II) chloride tetrahydrate (MnCl2 x 4H2O, *Sigma Aldrich*) was dissolved in Tris buffer at a dosage of 60 mg/kg for each animal injected into an osmotic mini-pump (*Alzet*) of 100µl volume. Pumps were primed in a water bath at 37°C overnight before surgical implantation to ensure their activation. Vetalgin (200mg/kg) was given subcutaneously as an analgesic 30 minutes prior to the surgery and mice were anesthetized with 4% isoflurane and secured to a stereotaxic frame. Immediately prior to the surgery, Meloxicam (0.5 mg/kg) was administered as an analgesic and eye ointment was applied to prevent drying of the eyes. Pumps were implanted subcutaneously under the skin of the left flank of the animal, while maintained under anesthesia with 2% isoflurane. Animals received Metacam in drinking water 3 days post-operation and were monitored for 7 days prior to imaging.

#### Magnetic resonance imaging

Manganese-enhanced MRI (MEMRI) was performed on a 9.4 T Bruker Biospec 94/20 system using a cross-coil configuration: radio-frequency transmission was achieved with a volume resonator, and signal detection was performed using room temperature 2 × 2 surface array coils. Six short consecutive 3D T1-weighted FLASH images were acquired with the following parameters: TR = 20.1 ms, TE = 5.86 ms, flip angle = 30°, matrix size = 200 × 150 × 100 voxels, field of view = 20 × 15 × 10 mm³, bandwidth = 25 kHz, two averages, and RF spoiling. Images were acquired along the rostral–caudal axis, with each 3D acquisition lasting 10 min 50 s. The total imaging session, including localizer scans and system adjustments, required approximately 75 minutes.

Data were converted using the Python Bru2nii function, in which the size of the voxels in the images was increased by tenfold, in order to be comparable with default values for human sized brains used by SPM12 in the following analyses. The six individual runs were coregistered, and a mean image calculated.

#### Image preprocessing

As the MEMRI data were acquired with a surface coil, correction of intensity inhomogeneities was essential. To ensure that bias field estimation was restricted to brain tissue, we generated a brain mask in native space. Because strong Mn²⁺ accumulation occurs in specific regions (e.g., cerebellum, olfactory bulb, and colliculi) and may distort bias field estimation, these regions were excluded from the mask.

The preprocessing pipeline was implemented in two passes. In the first round, images were only coarsely bias corrected, and the resulting masks were used to refine the second round of corrections. Preprocessing steps were performed using custom scripts and MATLAB R2024b and SPM12 (www.fil.ion.ucl.ac.uk/spm), if not otherwise stated.

a. Native mean MEMRI images were first coregistered to Hikishima space, followed by N4 bias correction (using 3D Slicer (vers. 5.8.1, www.slicer.org)) without a mask.
b. A preliminary brain mask was generated by segmenting the MEMRI images in SPM with the Hikishima template, summing the GM and WM compartments, and binarizing the result.
c. Individual masks were combined across subjects to create a group mask. Voxels were retained if they were included in at least two subjects.
d. This group mask was applied for brain extraction. A second segmentation was then performed using modified Hikishima templates in which ventricular and cortical CSF were separated, and brain extraction was repeated with this refined mask.
e. Brain-extracted images were normalized to the Allen Brain Atlas (ABA; AVGT.nii) using the “old normalization” function in SPM12.
f. The deformation fields were inverted to enable registration of ABA images into individual native space. Using this approach, we generated subject-specific ABA masks excluding the cerebellum, olfactory bulb, and inferior colliculi (defined via their ROI identifiers in ANO.nii).

In the second run of the pipeline, the original MEMRI images were bias corrected again (3D Slicer), this time using the refined ABA-derived brain masks in subject’s native space. Steps (d) and (e) were then repeated on these newly corrected images. Finally, data were smoothed using a kernel of 2 mm.

### c-Fos mapping

#### Tissue extraction for c-Fos

c-Fos expression was induced before sacrifice using the aforementioned open field & social interaction paradigm as a mild social challenge 60 minutes prior to perfusion. Animals (n = 32; divided equally by developmental stress treatment and sex) were then anesthetized with isoflurane and perfused trans-cardially with 4% paraformaldehyde (PFA). Brains were dissected and fixed overnight in PFA, then for the following 48 hours in phosphate-buffered saline (PBS)-30% sucrose, and then in PBS at 4°C.

#### Brain sectioning

Whole brains were placed in a container, bathed with optimal cutting temperature compound (OCT) and frozen, then stored at −80°C. Brains were sliced at 40um using a cryotome *(LEICA CM1900*) and stored in PBS or antifreeze solution.

#### Immunohistochemistry

In order to maintain specificity of c-Fos detection, tissue slices underwent blocking and permeabilization using bovine serum albumin and Triton X-100 respectively. Slices were then incubated with recombinant rabbit anti-c-Fos as a primary antibody to bind to the c-Fos protein, and donkey anti-rabbit Alexa Fluor-647 (AF647) as a secondary antibody to bind to the primary and therefore mark c-Fos molecules with a red fluorescent tag. Following incubation, slices were mounted onto slides and covered with DAPI, a nuclear stain.

#### Image acquisition and analysis

Images were acquired with slide scanner system Axioscan by Zeiss, using channels for DAPI and AF647. The tool Aligning Big Brain Atlases (ABBA)^45^ was used to align images to the Allen Brain Atlas in order to have an accurate per-region quantification of c-Fos-expressing neurons. BraiAn’s extension for QuPath^46^ v0.4.4 was used to segment c-Fos+ cells in the aligned images, as well as to localize image artifacts.

### Bulk RNA sequencing

#### Sacrifice

After behavioral testing was finished, animals were left undisturbed for over 24 hours before being sacrificed under baseline conditions. A subset of animals, balanced for sex and developmental condition, underwent restraint stress 4 hours prior to sacrifice. Animals were anesthetized with isoflurane and sacrificed by decapitation. Brains were dissected out and snap-frozen using methyl-butane on dry ice, and stored at −80°C.

#### Sample preparation and sequencing

Frozen brains were sectioned using a cryostat microtome at 250uM and a 0.8mm-diameter punching needle was used to extract tissue from the following brain regions: 1. prelimbic and 2. infralimbic prefrontal cortex (pPFC & iPFC), 3. nucleus accumbens (NAc), 4. anterodorsal bed nucleus of the stria terminalis (BNST), 5. paraventricular nucleus of the hypothalamus (PVN), 6. basolateral amygdala (BLA), 7. dorsal dentate gyrus and 8. dorsal CA1 of the hippocampus (dDG & dCA1), and 9. ventral dentate gyrus and 10. ventral CA1 (vDG & vCA1).

Prior to preparation, samples were randomized to reduce experimental batches using R package Omixer and subdivided into 5 plates each, including 2 brain regions; each brain region was treated as an individual experiment. RNA isolation was performed using the Zymo Direct-Zol kit and quality measured using the Agilent Tape Station 2000 (Supplementary Table 5), small RNA kit and Bioanalyzer, Agilent RNA ScreenTape and High Sensitivity RNA ScreenTape. cDNA libraries were prepared using the NEBNext Ultra II Direction RNA Library Prep Kit for Illumina, and sequencing was preformed using a NovaSeq6000 and Helmholtz Munich in two separate runs, using an S4 flow cell for the first four plates and S1 for the fifth plate.

### Statistical analysis

#### DeepOF analysis

Videos were analyzed using automated pose estimation software DeepLabCut v2.2.2^28^, and annotated motion tracking program DeepOF v0.6.4 and v07.0.2^29^. Behavioral outcomes were categorized using principal component analysis (PCA) to reduce dimensionality from all individual behaviors social and anxiety-related behaviors output by the DeepOF package.

#### Social box analysis

Interactions between mice were analyzed using DeepLabCut tracking and a previously published pipeline^47^, which calculated a David Score as a metric based on chasing behavior as a readout of social dominance rank.

#### Behavioral phenotyping statistics

For behavioral tests, R version 4.3.3 was used to assess statistical significance and create plots. For OFSI and FED3, normality was assessed with Shapiro-Wilk tests, and homogeneity of variance by Levene’s tests. When assumptions of normality and homogeneity were met, ANOVA was used for one, two, or three-variable designs. Significant ANOVAs were followed up by post-hoc Tukey tests for mean comparisons. A repeated-measured ANOVA was used to compare experimental groups when measurements were taken from multiple timepoints, and a mixed effects model was implemented in case of any missing values. Significance was set at *p*<0.05 and outliers were determined via the interquartile range method. For the social box, ordinal logistic regression was used to model the effect of stress on rank in the social dominance hierarchy.

#### MEMRI analysis

Statistical analysis was performed in SPM12, using a 2 x 2 factorial design with factors “sex” and “stress”, assuming independence and unequal variance between groups (male, ctrl: n=9; male, PNELS: n=8; female, ctrl: n=12; female, PNELS: n=11). As an explicit mask, we used the ABA mask image excluding the cerebellum and the olfactory bulb. Global normalization was considered by introducing global signal intensity as a nuisance regressor in the model. Statistical maps are presented with clusters surviving FDR cluster correction (p_FDR,cluster_ < 0.05), using a collection threshold of uncorrected p < 0.005.

#### c-Fos mapping analysis

For the whole-brain c-Fos analysis, mean-centered task partial least squares analysis (PLS) was conducted in python with BraiAn^45^. The analysis was done over 176 “summary structures” (a list of regions defined by CCFv3, Wang Q. et al. 2020), within Isocortex, CTXsp, HPF, STR, PAL, TH and HY. The PLS was performed over the c-Fos density of each brain structure, and the resulting contrasts and salience scores were generalized and normalized through permutation and bootstrap testing sampled 10,000 times each. In order to circumvent possible biases introduced by performing the immunochemistry staining at different time points—thus, with different solutions, we performed a batch normalisation. All region-quantifications were normalised on an additional staining that was performed the same day with the same solution throughout all 32 brains, 5 sections each.

PLS is presented for regions where p < 0.05 for the comparison between ctrl and PNELS for each sex. For connectivity analysis, Pearson correlation was calculated for cFos density across regions and presented for correlations where R ≥ 0.87 and p ≤ 0.0025. Linear regression was used to predict effects of relative density of cFos for individual brain regions of interest on the behavioral composite. Regions were chosen from those selected for bulk RNA sequencing due to known stress-related roles. Linear model formulas were formatted as: lm(region ∼ sex * treatment * DeepOF PC1).

#### Bulk RNA-seq analysis

For RNA sequencing, raw data was processed using the nf-core bioinformatic pipeline *rnaseq* (version 3.5; Ewels et al., 2020), in which *STAR* was used for alignment to the reference genome mm10 for mouse (refGenie), and *salmon* for gene quantification. Samples with an alignment rate of less than 5% were excluded and gene expression was filtered to a minimum of 10 counts for subsequent analysis. Principal component analysis was used to further identify outliers and exclude 36 of the 469 samples.

Differential gene expression was used to define transcripts significantly altered. DEGs were determined using the DESeq2 R package, with a sex-stratified approach using likelihood ratio tests, which captured the individual effects of PNELS and adulthood stress, as well as the interaction between them. The threshold used to define differential expression was a padj < 0.1 Full DESeq2 results are found in Supplementary Table(s) XYZ.

Rank-Rank hypergeometric overlap (RRHO) analysis, a threshold-free method to compare genetic signatures across studies^48^, was used to compare transcriptomic profiles across studies. RRHO was conducted using the RRHO2 R package. Lists were filtered to include only genes overlapping between studies and the significance was mapped using -log10(padj) adjusted for direction of change in expression, using data from Labonté and colleagues^33^ for the interspecies comparison. Fisher’s exact tests were used to assess the significance of overlapping differentially expressed genes (DEGs) between datasets for each quadrant of concordant and discordant regulation, using the intersection of tested genes as background. For the sex-specific comparisons, global concordance and discordance of gene expression changes between males and females were quantified using the RRHO correlation (ρ), the proportion of genes changing in the same direction (sign concordance), and 95% confidence intervals.

Gene ontology enrichment analysis was performed with the R package clusterProfiler. For mouse RRHO, discordant genes in the BNST and NAc were set against a background of all genes overlapping between sexes for each region for gene ontology enrichment (enrichGO, cutoff = 0.1). For the interspecies comparison in the nucleus accumbens, gene set enrichment analysis (gseGO) was used to account for significance and directionality (p cutoff = 0.05). R package GoSemSim was used to determine semantic similarity between significantly enriched (p < 0.05) human and mouse GO terms from NAc data. Dotplots include top human terms with highest similarity to a mouse term (> 0.6), with the point color and size for each term determined by them mouse term’s p-value and enrichment score respectively.

Weighted gene co-expression network analysis^49^ via the WGCNA package was used to identify patterns of gene co-expression and determine functionally-relevant networks of genes in an unsupervised manner. Samples were stratified by region, as well as adulthood condition to avoid skewing of module detection. Pearson correlation of module eigengene (ME) values to traits was used to identify modules associated with PNELS, PC1 eigenvalues of DeepOF behaviors, and sex. For select modules with associations (R > 0.5, p < 0.1) to any of the traits, linear regression was used to assess how well sex, PNELS, and ME value predict the behavioral score.

## Acknowledgements

We thank Clara Engelhardt, Andrea Ressle, Cornelia Flachskamm, Daniela Harbich, Bianca Schmid, Lotte van Doeselaar, Benoit Boulat and Anthi Krontira for technical assistance. In addition, we would like to thank Linda Requie, Carmen Camarena, Alessandro Bellone, Caterina Giovannini, Alice Jelmoni, Marco Bressi, Stefania Sempreboni, Alessia Marchesin, Edoardo Cavaglià, and Maddalena Pieroni for their support in whole brain cFos mapping.

## Data and Materials Availability

All data needed to evaluate the conclusions in the paper are present in the paper and/or the Supplementary Materials. Additional data related to this paper may be requested from the authors. Processed data files are available upon request before publication and will be made publicly available upon publication.

## Supplementary

**Supplementary Figure 1.**
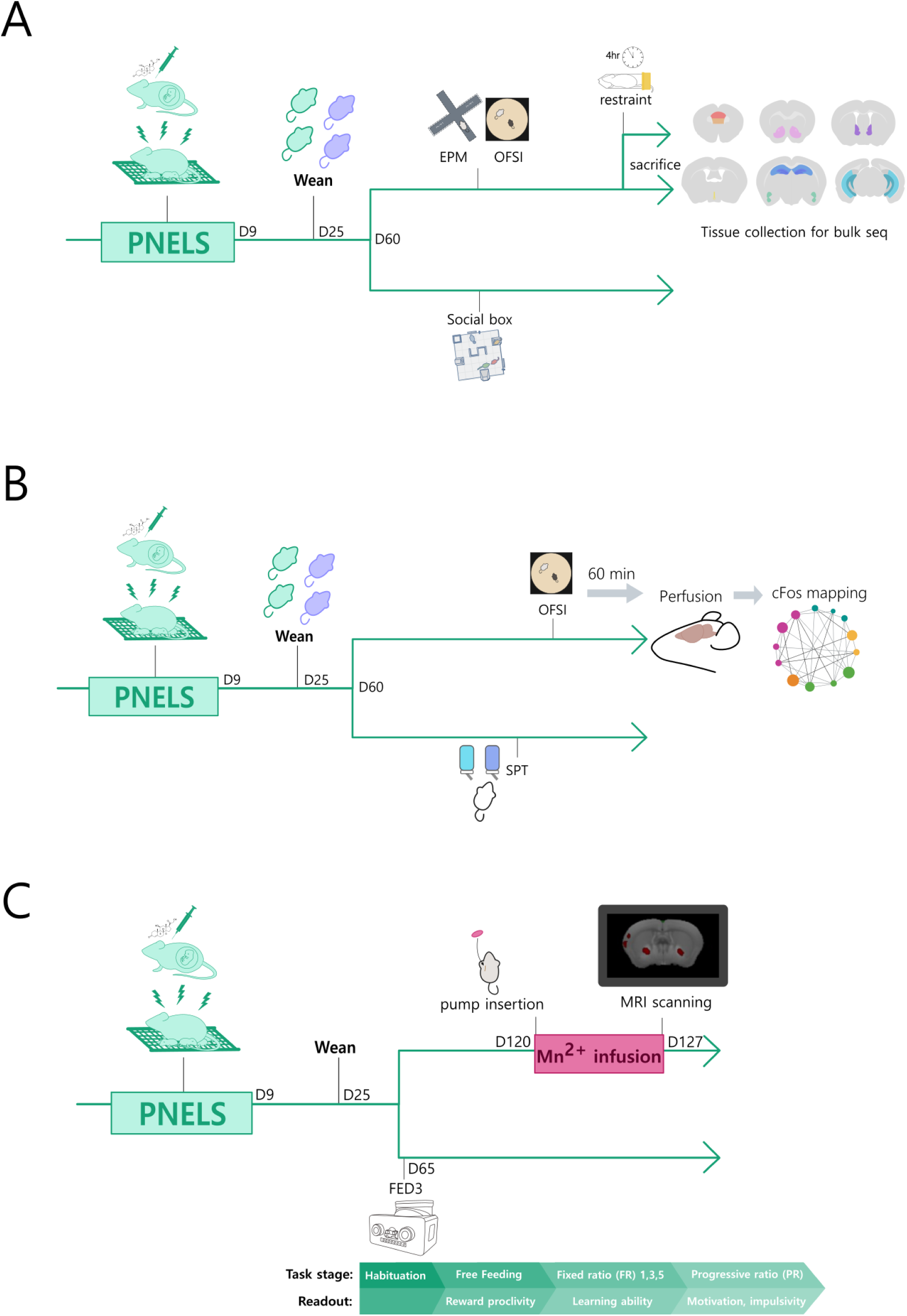
Cohort timelines. Experimental timelines for each cohort. Behavioral phenotyping and bulk RNA sequencing were done in the first cohort (**A**), behavior and cFos mapping in the second (**B**), and MRI in the 3^rd^ (**C**). Each timeline bifurcation represents a different subset of animals.

**Supplementary Figure 2.**
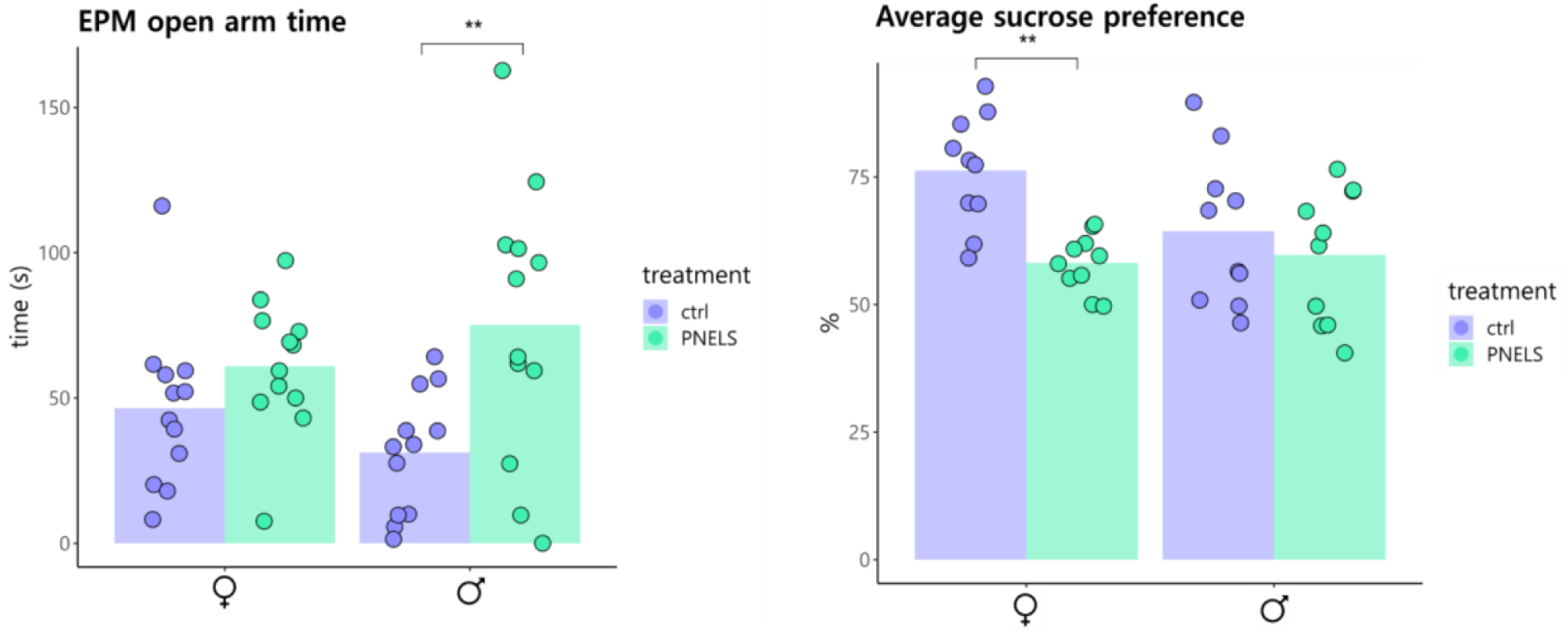
Classical behavioral assessments. Classical assessments of stress-related behaviors did not capture nuanced and sex specific effects of developmental stress (**EPM**: treatment: F(1, 44) = 10.093, p = 0.00272; sex: ns; sex:PNELS: ns; Post-hoc Tukey: M ctrl vs PNELS: p =0.00798. **SPT**: treatment: F(1, 36) = 9.526, p = 0.00388; sex: ns; sex:PNELS: ns; Post-hoc Tukey: F ctrl vs PNELS: p =0.00717).

**Supplementary Figure 3.**
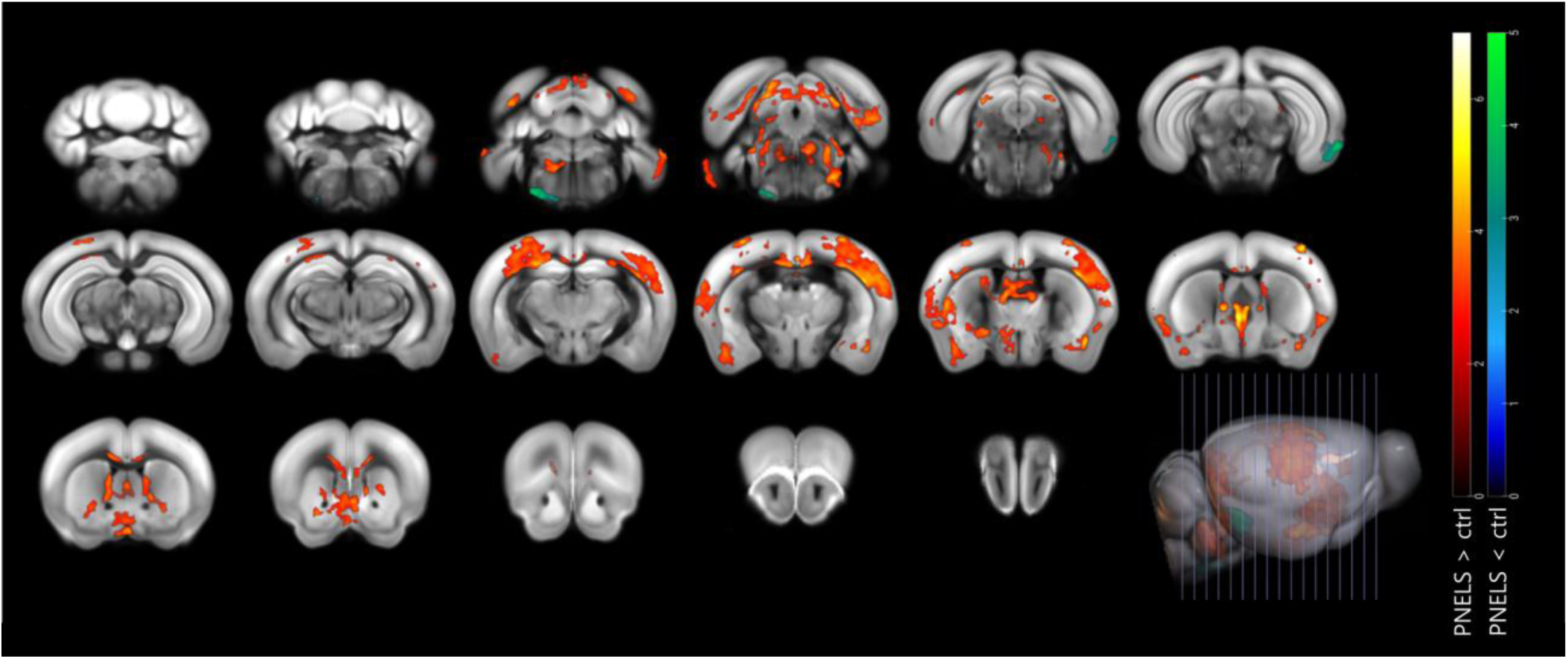
MEMRI stress effects. The main effect of stress complements interaction in many region, such as Superior Colliculus, Entorhinal areas, Lateral Septum, Medial Preoptic Area, cortices (primary somatosensory, Posterolateral visual area, Agranular Insula posterior part), Endopiriform nucleus ventral part, and central amygdala. Stress predominantly increased Mn2+ intensity. Statistical maps are presented with clusters surviving FDR cluster correction (pFDR,cluster < 0.05), using a collection threshold of uncorrected p < 0.005.

**Supplementary Figure 4.**
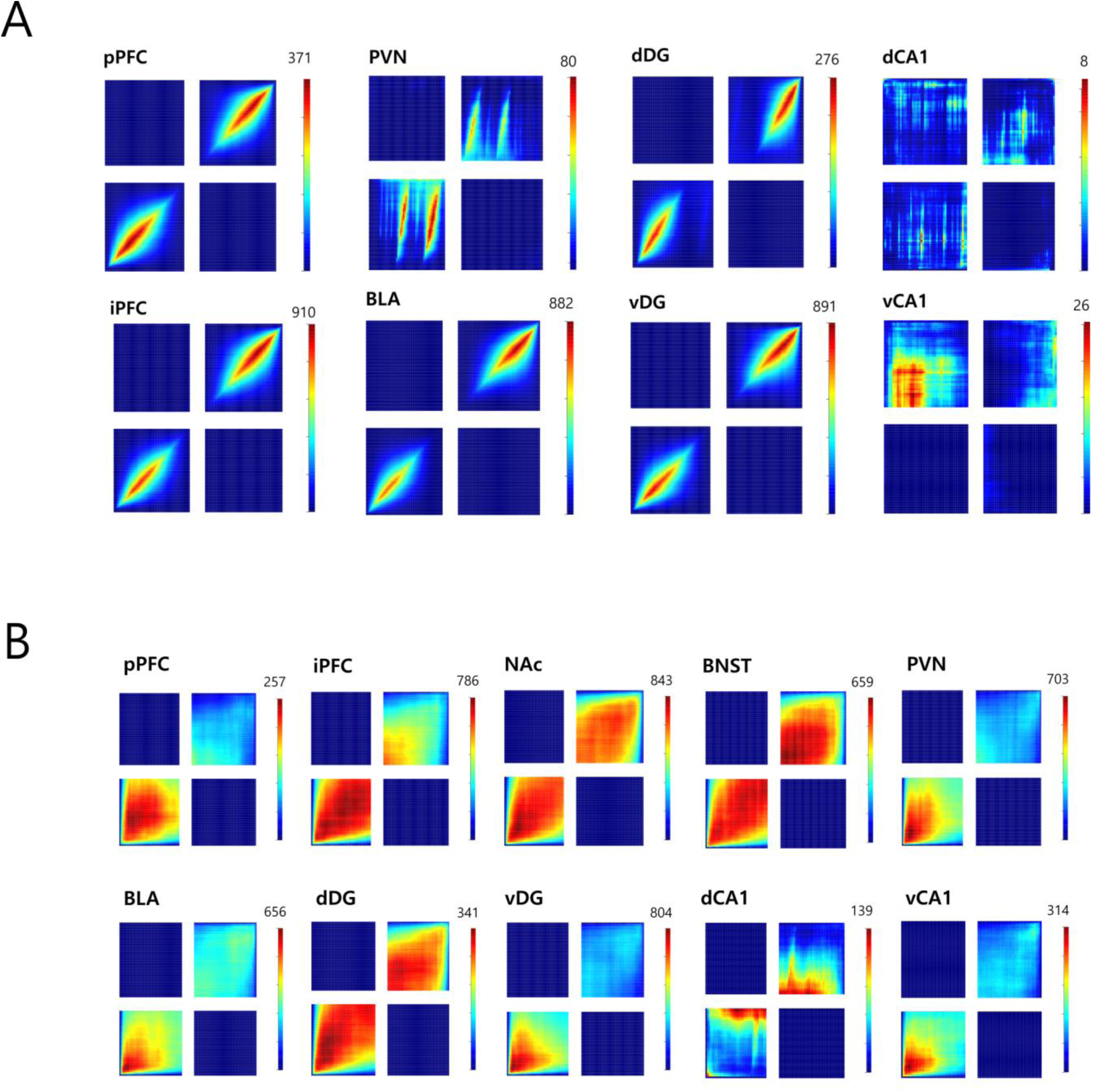
RRHO plots for all collected regions. RRHO comparing male and female PNELS results show mainly concordance of transcriptional signatures for all collected regions (**A**) except NAc and BNST (Fig. 4B). RRHO for restraint results show exclusive concordance between sexes, highlighting the unique sex-specificity associated with PNELS in select regions (**B**).

**Supplementary Figure 5.**
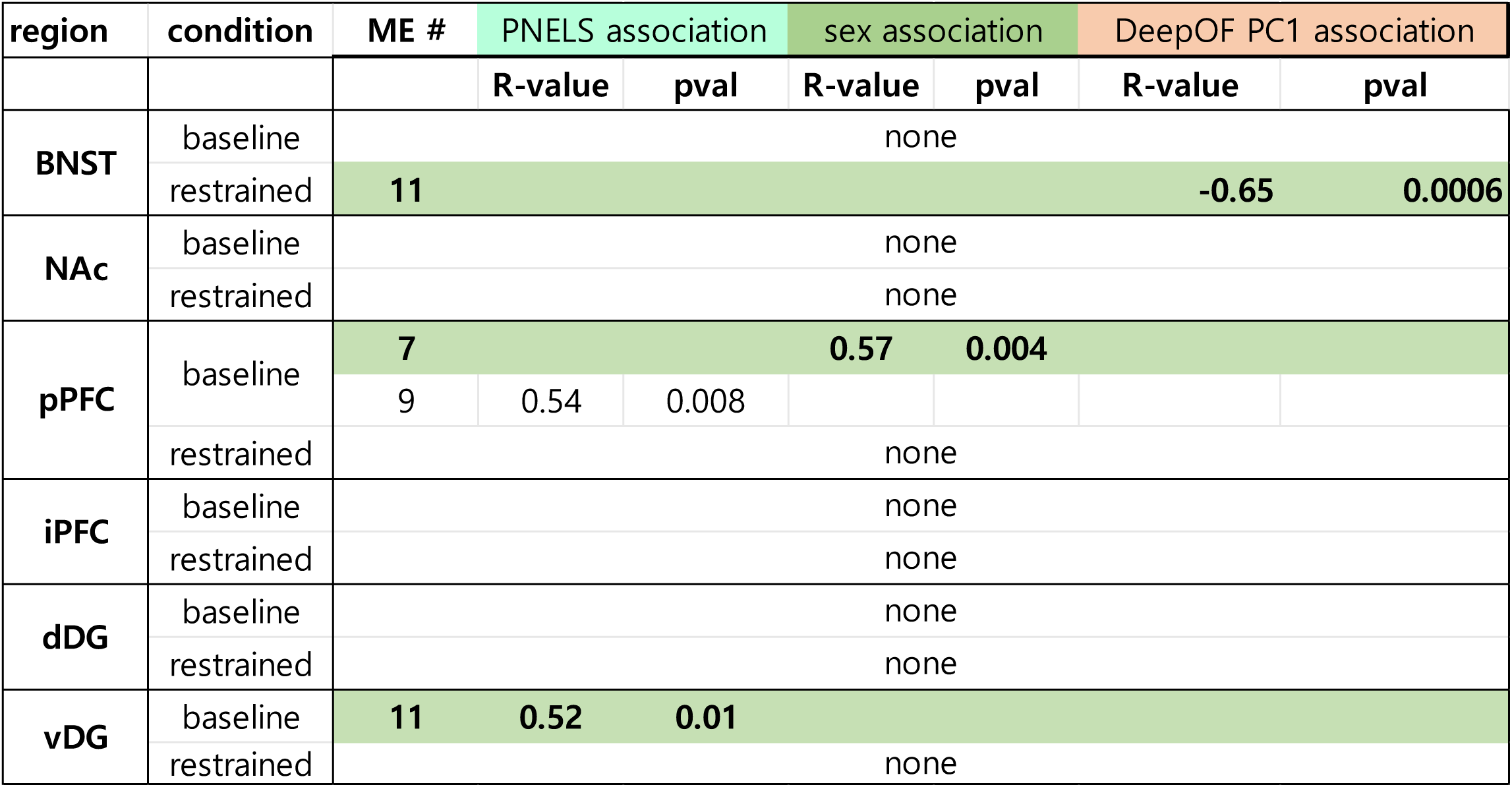
WGCNA modules. Modules found with strong and significant (R > 0.5, p < 0.05) associated to PNELS, sex, or DeepOF PC1 for regions chosen for WGCNA. Modules highlighted in green were analyzed for predictive effects on behavior (Fig. 4D).

## References

1. Kessler, R. C. et al. Childhood adversities and adult psychopathology in the WHO World Mental Health Surveys. Br. J. Psychiatry 197, 378–385 (2010).

2. Chapman, D. P. et al. Adverse childhood experiences and the risk of depressive disorders in adulthood. J. Affect. Disord. 82, 217–225 (2004).

3. Cohen, P., Brown, J. & Smaile, E. Child abuse and neglect and the development of mental disorders in the general population. Dev. Psychopathol. 13, 981–999 (2001).

4. McLaughlin, K. A. et al. Childhood adversities and adult psychopathology in the National Comorbidity Survey Replication (NCS-R) III: Associations with functional impairment related to DSM-IV disorders. Psychol. Med. 40, 847–859 (2010).

5. Beery, A. K. & Zucker, I. Sex bias in neuroscience and biomedical research. Neurosci. Biobehav. Rev. 35, 565–572 (2011).

6. Schraufnagel, T. J., Wagner, A. W., Miranda, J. & Roy-Byrne, P. P. Treating minority patients with depression and anxiety: what does the evidence tell us? Gen. Hosp. Psychiatry 28, 27–36 (2006).

7. Galea, L. A. & Parekh, R. S. Ending the neglect of women’s health in research. BMJ 381, p1303 (2023).

8. Rechlin, R. K., Splinter, T. F. L., Hodges, T. E., Albert, A. Y. & Galea, L. A. M. An analysis of neuroscience and psychiatry papers published from 2009 and 2019 outlines opportunities for increasing discovery of sex differences. Nat. Commun. 13, 2137 (2022).

9. Turano, A., Osborne, B. F. & Schwarz, J. M. Sexual Differentiation and Sex Differences in Neural Development. in Neuroendocrine Regulation of Behavior (eds. Coolen, L. M. & Grattan, D. R.) 69–110 (Springer International Publishing, Cham, 2019). doi:10.1007/7854_2018_56.

10. Benavides, A. et al. Sex-specific alterations in preterm brain. Pediatr. Res. 85, 55–62 (2019).

11. Oyola, M. G. & Handa, R. J. Hypothalamic-pituitary-adrenal and hypothalamic-pituitary-gonadal axes: sex differences in regulation of stress responsivity. Stress Amst. Neth. 20, 476–494 (2017).

12. Papadopoulos, A. D. & Wardlaw, S. L. Testosterone suppresses the response of the hypothalamic-pituitary-adrenal axis to interleukin-6. Neuroimmunomodulation 8, 39–44 (2000).

13. Brix, L. M. et al. Metabolic effects of early life stress and pre-pregnancy obesity are long lasting and sex specific in mice. Eur. J. Neurosci. 58, 2215–2231 (2023).

14. Bordes, J. et al. Sex-specific fear acquisition following early life stress is linked to amygdala and hippocampal purine and glutamate metabolism. *Commun*. Biol. 7, 1–12 (2024).

15. van Doeselaar, L. et al. FKBP51 in glutamatergic forebrain neurons promotes early life stress inoculation in female mice. Nat. Commun. 16, 2529 (2025).

16. Guadagno, A., Belliveau, C., Mechawar, N. & Walker, C.-D. Effects of Early Life Stress on the Developing Basolateral Amygdala-Prefrontal Cortex Circuit: The Emerging Role of Local Inhibition and Perineuronal Nets. Front. Hum. Neurosci. 15, 669120 (2021).

17. Kronman, H. et al. Long-term behavioral and cell-type-specific molecular effects of early life stress are mediated by H3K79me2 dynamics in medium spiny neurons. Nat. Neurosci. 24, 667–676 (2021).

18. Weinstock, M. Sex-dependent changes induced by prenatal stress in cortical and hippocampal morphology and behaviour in rats: an update. Stress Amst. Neth. 14, 604–613 (2011).

19. Wu, Y., De Asis-Cruz, J. & Limperopoulos, C. Brain structural and functional outcomes in the offspring of women experiencing psychological distress during pregnancy. Mol. Psychiatry 29, 2223–2240 (2024).

20. Sutherland, S. & Brunwasser, S. M. Sex Differences in Vulnerability to Prenatal Stress: A Review of the Recent Literature. Curr. Psychiatry Rep. 20, 102 (2018).

21. Nold, V. et al. Impact of Fkbp5 × early life adversity × sex in humanised mice on multidimensional stress responses and circadian rhythmicity. Mol. Psychiatry 27, 3544–3555 (2022).

22. Bangasser, D. A. & Cuarenta, A. Sex differences in anxiety and depression: circuits and mechanisms. Nat. Rev. Neurosci. 22, 674–684 (2021).

23. Baugher, B. J. et al. Sub-chronic stress induces similar behavioral effects in male and female mice despite sex-specific molecular adaptations in the nucleus accumbens. Behav. Brain Res. 425, 113811 (2022).

24. McCarthy, M. M., Arnold, A. P., Ball, G. F., Blaustein, J. D. & De Vries, Geert. J. Sex Differences in the Brain: The Not So Inconvenient Truth. J. Neurosci. 32, 2241–2247 (2012).

25. Bangasser, D. A. & Wicks, B. Sex-specific mechanisms for responding to stress. J. Neurosci. Res. 95, 75–82 (2017).

26. Schmidt, M. V. The tricky business of predicting the future: How brain and body adapt to early life adversity. Neurosci. Biobehav. Rev. 155, 105466 (2023).

27. Musillo, C., Berry, A. & Cirulli, F. Prenatal psychological or metabolic stress increases the risk for psychiatric disorders: the “funnel effect” model. Neurosci. Biobehav. Rev. 136, 104624 (2022).

28. Mathis, A. et al. DeepLabCut: markerless pose estimation of user-defined body parts with deep learning. Nat. Neurosci. 21, 1281–1289 (2018).

29. Bordes, J. et al. Automatically annotated motion tracking identifies a distinct social behavioral profile following chronic social defeat stress. Nat. Commun. 14, 4319 (2023).

30. Miranda, L., Bordes, J., Pütz, B., Schmidt, M. V. & Müller-Myhsok, B. DeepOF: a Python package for supervised and unsupervised pattern recognition in mice motion tracking data. J. Open Source Softw. 8, 5394 (2023).

31. Forkosh, O. et al. Identity domains capture individual differences from across the behavioral repertoire. Nat. Neurosci. 22, 2023–2028 (2019).

32. Matikainen-Ankney, B. A. et al. An open-source device for measuring food intake and operant behavior in rodent home-cages. eLife 10, e66173 (2021).

33. Labonté, B. et al. Sex-specific transcriptional signatures in human depression. Nat. Med. 23, 1102–1111 (2017).

34. Stadtler, H. & Neigh, G. N. Sex Differences in the Neurobiology of Stress. Psychiatr. Clin. North Am. 46, 427–446 (2023).

35. Hodes, G. E., Bangasser, D., Sotiropoulos, I., Kokras, N. & Dalla, C. Sex Differences in Stress Response: Classical Mechanisms and Beyond. Curr. Neuropharmacol. 22, 475–494 (2024).

36. Rainville, J. R., Lipuma, T. & Hodes, G. E. Translating the Transcriptome: Sex Differences in the Mechanisms of Depression and Stress, Revisited. Biol. Psychiatry 91, 25–35 (2022).

37. Altemus, M., Sarvaiya, N. & Epperson, C. N. Sex differences in anxiety and depression clinical perspectives. Front. Neuroendocrinol. 35, 320–330 (2014).

38. Basar, K. et al. Nucleus accumbens and impulsivity. Prog. Neurobiol. 92, 533–557 (2010).

39. Galea, L. A. M. et al. Endocrine regulation of cognition and neuroplasticity: our pursuit to unveil the complex interaction between hormones, the brain, and behaviour. Can. J. Exp. Psychol. Rev. Can. Psychol. Exp. 62, 247–260 (2008).

40. Kos, A. et al. Early life adversity shapes social subordination and cell type–specific transcriptomic patterning in the ventral hippocampus. Sci. Adv. 9, eadj3793 (2023).

41. Russo, S. J. & Nestler, E. J. The brain reward circuitry in mood disorders. Nat. Rev. Neurosci. 14, 609–625 (2013).

42. Lebow, M. A. & Chen, A. Overshadowed by the amygdala: the bed nucleus of the stria terminalis emerges as key to psychiatric disorders. Mol. Psychiatry 21, 450–463 (2016).

43. Astiz, M. et al. The circadian phase of antenatal glucocorticoid treatment affects the risk of behavioral disorders. Nat. Commun. 11, 3593 (2020).

44. Rice, C. J., Sandman, C. A., Lenjavi, M. R. & Baram, T. Z. A novel mouse model for acute and long-lasting consequences of early life stress. Endocrinology 149, 4892–4900 (2008).

45. Chiaruttini, N. et al. ABBA+BraiAn, an integrated suite for whole-brain mapping, reveals brain-wide differences in immediate-early genes induction upon learning. Cell Rep. 44, 115876 (2025).

46. Bankhead, P. et al. QuPath: Open source software for digital pathology image analysis. Sci. Rep. 7, 16878 (2017).

47. Karamihalev, S. et al. Social dominance mediates behavioral adaptation to chronic stress in a sex-specific manner. eLife 9, e58723 (2020).

48. Plaisier, S. B., Taschereau, R., Wong, J. A. & Graeber, T. G. Rank–rank hypergeometric overlap: identification of statistically significant overlap between gene-expression signatures. Nucleic Acids Res. 38, e169 (2010).

49. Langfelder, P. & Horvath, S. WGCNA: an R package for weighted correlation network analysis. BMC Bioinformatics 9, 559 (2008).

